## Supplemental Text, Table, and Figures for "Immune Lag Is a Major Cost of Prokaryotic Adaptive Immunity During Viral Outbreaks"

June 29, 2021

#### SText 1 Model Details

##### SText 1.1 Chemostat Assumption

Our model follows the chemostat formalism that has been standard in models of virus-microbe systems for decades [20, 6, 12, 24]. Under this setup, media flows into the system at some set rate, $w$, from a nutrient reservoir with resource concentration $r_0$. Media is removed from the system at the same constant rate $w$ in order to maintain a constant volume. Cell and virion populations are expressed as concentrations, and cell and viral particle volumes are neglected for simplicity. Chemostat setups are useful in that their continuous (i.e., not serial dilution) and well-mixed (i.e., no spatial structure) nature makes them analytically tractable and easy to work with mathematically, while still being conducive to experimentation in the lab (i.e., the modeled system corresponds well to a realistic laboratory setup). Additionally, the chemostat assumption of constant resource inflow may correspond well to some natural systems (at least over shorter timescales), and has been applied previously to models of natural systems (e.g., [25]). Our modification to create a "virostat" maintains these desirable features, and simply adds a set concentration of viral particles, $v_0$ to the media reservoir. In doing so, we also bring our model closer to natural systems where viral immigration is likely to pose a constant challenge for microbes.

##### SText 1.2 Resource Uptake

Because we model resources explicitly as part of our chemostat model, we need to also model realistic resource uptake dynamics. We follow standard Monod growth kinetics for resource uptake, with half-saturation constant $z$ [17]. We assume that both immune and susceptible host have similar resource uptake kinetics, since it would be unlikely that the acquisition of a CRISPR spacer would lead to changes in growth parameters. For the SM strain, we assume that the loss of the phage receptor decreases the maximal growth rate, $v$, of the population by some set cost ($0 \leq \kappa \leq 1$). In principle, the loss of a receptor could also impact the half-saturation constant, but we do not consider this scenario here. Note that in our most simplified lag model (see S4 Text) we model resources implicitly as logistic growth, and that this change in model setup does not alter our qualitative results.

#### SText 1.3 Virus-Host Dynamics

Virus-host dynamics in our model follow an obligately-lytic viral lifestyle. The burst size and latency periods are assumed to remain constant. Some models allow burst size to vary with nutrient availability, but for the sake of simplicity we do not consider these more complicated forms here. We implicitly model latency period by incorporating an infected class ($I$) into our model. In effect, this model structure assumes that the latency period of an infection is exponentially distributed with mean $1/\gamma$, and this assumption and corresponding model structure is widely used in the literature. Alternatively, it is possible to explicitly model a fixed latency period by using delay differential equations, but the analysis of such models is much more difficult and coming up with a good set of initial conditions for such models is non-trivial [24]. As such, we prefer the simpler and more tractable implicit approach. Finally, we do not model viral decay in our model as we assume the flow of viral particles out of the system is much faster than their decay times.

#### SText 1.4 CRISPR Spacer Acquisition

Our model assumes that only a small number of infections will lead to immunization. That is, some small fraction of infections ($\mu$) results in the acquisition of a spacer, whereas the rest $(1-\mu)$ result in death of the host and successful reproduction of the virus. This assumption is consistent both with previous models and with experimental evidence that shows that CRISPR spacer acquisition events are quite rare [10, 11, 4, 16, 22]. Additionally, we neglect any coevolutionary dynamics in this simple model (this assumption is relaxed during simulations of repeated outbreaks), so that a single spacer acquisition even is sufficient to completely prevent viral infection and no viral escape mutants arise. Because we do not model viral escape, it is not necessary to model the length of the CRISPR array or the diversity of spacers in the overall population, since the only relevant distinction will be between hosts with no spacers and hosts with at least one spacer (similar to the analysis in [4]). While coevolutionary dynamics are interesting, they are beyond the scope of the current study. We show that even in the best-case scenario for CRISPR (i.e., no possibility for viruses to escape targeting), immune lag can still lead to situations where CRISPR is a sub-optimal defense strategy. The addition of coevolutionary dynamics would only serve to even more strongly disfavor the evolution of a CRISPR defense strategy.

### SText 2 Autoimmunity is Unlikely to Lead to the Invasion of Alternative Costly Defense Strategies to CRISPR-Cas

In order to quantify the effects of autoimmunity on CRISPR-encoding host we explicitly modeled CRISPR-Cas's inherent capacity for self versus non-self recognition in our model by developing a replicon-based model of spacer acquisition that takes into account the demonstrated preference of the Cas acquisition machinery for double-strand breaks occurring at collapsed replication forks [13]. We consider the following modification of the model in the main text:

$$
\begin{aligned}
\overbrace{\dot{R}}^{\text{Resources}} &= \overbrace{w(r_0 - R)}^{\text{Flow}} - \overbrace{\frac{evR}{z+R}(S+C)}^{\text{Resource Uptake by Cells}} \\
\overbrace{\dot{S}}^{\text{Susceptible}} &= \left( \overbrace{\frac{vR}{z+R}}^{\text{Growth}} - \overbrace{\delta V}^{\text{Infection}} - \overbrace{w}^{\text{Flow}} - \overbrace{2\mu_r P}^{\text{Autoimmunity}} \right) S \\
\overbrace{\dot{C}}^{\text{Immune}} &= \left( \overbrace{\frac{vR}{z+R}}^{\text{Growth}} - \overbrace{w}^{\text{Flow}} - \overbrace{2\mu_r P}^{\text{Autoimmunity}} \right) C + \overbrace{\mu_r I_v}^{\text{Immunization}} \\
\overbrace{\dot{I}}^{\text{Infected}} &= \overbrace{\delta SV}^{\text{Infection}} - \overbrace{\gamma I}^{\text{Lysis}} - \overbrace{wI}^{\text{Flow}} - \overbrace{\mu_r I_v}^{\text{Immunization}} \\
\overbrace{\dot{I}_v}^{\text{Intracellular Virions}} &= \overbrace{\delta SV}^{\text{Infection}} + \overbrace{\gamma \log(\beta) I_v}^{\text{Reproduction}} - \overbrace{\left( \frac{\gamma e^{\xi \log \beta}}{\xi} \right) I}^{\text{Immunization Window}} - \overbrace{wI_v}^{\text{Flow}} - \overbrace{\mu_r \left( \frac{I_v^2}{I} \right)}^{\text{Immunization}} \\
\overbrace{\dot{V}}^{\text{Viruses}} &= \overbrace{w(v_0 - V)}^{\text{Flow}} + \overbrace{\beta \gamma I}^{\text{Lysis}} - \overbrace{\delta (S+C) V}^{\text{Adsorption}} .
\end{aligned}
\tag{1}
$$

where we track the number of intracellular viral replicons from which spacers can be acquired ($I_v$), which is a significant departure from previous models. We model the immunization process as being specifically dependent on the number of viral replicons in a cell ($\mu_r$ is the per-replicon/time rate of spacer acquisition, see discussion below), assuming the majority of spacers are acquired from collapsed replication forks [13]. We assumed exponential growth of the intracellular viral population and defined a time-window ($\xi$) in which immunization is possible (accounting for the fact that spacer acquisition likely happens early on during infection [16]; changing the model to linear viral production makes little difference in our conclusions, see below).

In addition to immunization, the process of auto-immunization will also be replicon-based. Bacterial cells may have high "effective ploidy" [5, 15], where cells require multiple active replication centers to achieve high growth rates given physical limitations on the rate of DNA replication [1, 18]. Let $P$ be the minimum number of simultaneously occurring DNA replication cycles required to observe a given host growth rate and let $\alpha$ be the maximum rate of genome replication on a single strand (theoretically $\sim$ 600 bp/sec [27, 24]). Assuming that $\alpha$ is constant across growth rates, $P$ can be expressed at the rate of host cell duplication divided by the rate of host genome duplication:

$$
P = \frac{vR}{\alpha(z+R)}. \tag{2}
$$

Thus, assuming bidirectional replication, and letting $\mu_r$ be the per-replicon-per-hour spacer acquisition rate, the rate at which self targeting spacers are acquired will be $2\mu_r P$. Note that at low growth rates $P$ can drop below one, corresponding to the proportion of time that the genome must be actively replicating to obtain the observed growth rate. In reality bacteria may have multiple ways of further differentiating self from non-self [23] (e.g., by the presence of Chi sites on the host genome [13, 16]) so that our model represents a "worst case scenario" for the host in terms of autoimmunity.

In agreement with experimental data that detects no constitutive cost of CRISPR-Cas immunity [26, 2, 14], our model's equilibria indicated that there should be essentially no impact of autoimmunity on the hosts' fitness at realistic rates of spacer acquisition. In the presence of virus at equilibrium (enforced by letting $v_0 > 0$, though this condition is not necessary for this equilibrium to occur), all host will ultimately be defended (since undefended host also pay the cost of autoimmunity and there is no spacer loss in this model). The equilibrium state with host and virus present can be described as

$$\tilde{R} = \frac{zw\alpha}{v\alpha - 2v\mu_r - w\alpha}, \tag{3}$$

$$\tilde{D} = \frac{w(r_0 - \tilde{R})(z + \tilde{R})}{ev\tilde{R}}, \tag{4}$$

$$\tilde{V} = \frac{wv_0}{w + \delta\tilde{D}}, \tag{5}$$

where the appropriate substitutions for $\tilde{R}$ and $\tilde{D}$ can be made in equations 4 and 5 respectively (left in this form for simplicity). Note that $\tilde{S} = 0$, $\tilde{I} = 0$, and $\tilde{I_v} = 0$.

We found that a costly SM strategy was unable to invade a resident CRISPR-immune host population at equilibrium with realistic levels of autoimmunity. Suppose that we add a CRISPR-lacking surface-mutant (hereafter "SM"; $M$) to our system at equilibrium:

$$\dot{M} = \left(\frac{(1-\kappa)vR}{z + R} - w\right) M. \tag{6}$$

Observe that the SM population is immune to viral infection but experiences some fixed growth cost ($\kappa$). Can this mutant invade the population at equilibrium? The condition for the initial invasion of this mutant (when $M(0) \approx 0$) is that

$$\frac{(1-\kappa)v\tilde{R}}{z + \tilde{R}} - w > 0 \tag{7}$$

where $\tilde{R}$ is the resource concentration at equilibrium. This invasion condition can be re-written as

$$\frac{2\mu_r}{\alpha} > \kappa. \tag{8}$$

As a point of reference, in one popular type II-A model system there are around $10^{-7}$ immunizations per infection [16], which we can convert to an acquisition rate of $\mu \approx 10^{-6}$ spacers acquired per replicon per hour (see below). Assuming a host genome length of 5-6mb, we expect a maximum genome replication rate of $\alpha \approx 0.4$ per hour [27, 24].

Thus, given realistic parameter estimates, $\kappa$ must be exceedingly small for an SM mutant to invade (SS2a Fig). This is true even allowing for several orders of magnitude of error in our estimate of the spacer acquisition rate (S2a Fig). Yet, experiments show time and again that SM strategies do evolve, and often out-compete CRISPR-encoding host. Thus either the cost of receptor loss must be very small ($\kappa < 10^{-4}$) or autoimmunity is not the major factor in determining whether an SM mutant invades. Importantly, host Chi sites constrain the amount of self-targeting during CRISPR-Cas immunization [13], meaning that our estimate for the maximum cost of an SM strategy ($\kappa$)

that permits invasion is almost certainly an overestimate (i.e., the effects of autoimmunity are likely even less important for the evolution of host defense strategy than indicated by our model).

How high would rates of self-targeting need to be for autoimmunity to cause the invasion of an alternative SM strategy into a CRISPR-immune population? If we assume a moderate, 5% growth cost of an SM strategy, our per-replicon spacer acquisition rate would need to be at least 0.01 acquisitions per replicon per hour for that SM population to invade. This is many orders of magnitude above what we observe even in experimental systems specifically chosen for their ability to rapidly acquire spacers (e.g., $\mu_r \approx 10^{-6}$, see Sfigure S2 Figure; [3, 16]). Certainly we know that autoimmunity happens often enough to be detected in genomic datasets [19, 21], but it is unlikely to have much impact on host population and evolutionary dynamics [14, 2].

#### Derivation of Intracellular Virus Model

Consider the population of intracellular viral replicons ($I_v$):

$$\dot{I}_v = \overbrace{\delta UV}^{\text{Infection}} + \overbrace{\gamma \log(\beta) I_v}^{\text{Reproduction}} - \overbrace{\beta\gamma I}^{\text{Lysis}} - \overbrace{w I_v}^{\text{Washout}} - \overbrace{\mu_r \left(\frac{I_v^2}{I}\right)}^{\text{Immunization}} . \tag{9}$$

This equation requires some explanation. The expected rate of immunization, again assuming that spacer acquisition happens on a per-replicon basis, is

$$\overbrace{\mu_r \times I \times E[\text{viral replicons per cell}]}^{\text{Immunization}} = \mu_r I \left(\frac{I_v}{I}\right) = \mu_r I_v. \tag{10}$$

If infected cells lyse after an expected infection length $\frac{1}{\gamma}$, with burst size $\beta$, then the average rate of viral replication within these cells is $\gamma \log(\beta)$, assuming exponential increase after a single viral injection. We multiply the rate of infected cell washout ($wI$) by the expected replicons per cell ($\frac{I_v}{I}$) to obtain the rate of replicon washout ($wI_v$). Similarly, we multiply the rate of host immunization ($\mu_r I_v$) by the expected number of replicons lost during a single immunization even ($\frac{I_v}{I}$) to get the rate of replicon loss due to immunization ($\mu_r \left(\frac{I_v^2}{I}\right)$). The reader may observe that this model allows spacer acquisition to occur any time during the course of an infection, which is unlikely to say the least [16]. Thus, we modify equation 9 to allow for shorter windows of immunization with the addition of a parameter $\xi$, representing the proportion time during an infection in which spacer acquisition is possible, so that:

$$\dot{I}_v = \overbrace{\delta UV}^{\text{Infection}} + \overbrace{\gamma \log(\beta) I_v}^{\text{Reproduction}} - \overbrace{\left(\frac{\gamma e^{\xi \log \beta}}{\xi}\right) I}^{\text{Immunization Window}} - \overbrace{w I_v}^{\text{Washout}} - \overbrace{\mu_r \left(\frac{I_v^2}{I}\right)}^{\text{Immunization}} . \tag{11}$$

The third term in this equation can be found by considering that viruses within a cell undergo exponential growth at a rate $r = \gamma \log(\beta)$ starting from an initial population of a single virion (neglecting the possibility of multiple infection). Thus, in a single infected cell the number of replicating viruses ($V_i$) will be

$$V_i(t) = e^{\gamma \log(\beta) t}. \tag{12}$$

If the immunization window is of average length $\frac{\xi}{\gamma}$, then $V_i = e^{\xi \log \beta}$ at the end of this window and the rate at which cells leave this window will be $\frac{\gamma}{\xi}$.

#### Estimating The Spacer Acquisition Rate Per-Replicon

We base our rate estimate on a commonly used model system for CRISPR-Cas research: Staphylococcus aureus background expressing a type II-A CRISPR locus from Streptococcus pyogenes and infected by phage $\phi NM4\gamma4$. This phage has a burst size of approximately 80 virions/infection [9] and the time from infection to lysis is $\sim 45$ minutes (so that $\gamma = \frac{4}{3}$ lyses/hr [16]). Approximately 1 in $10^7$ infections lead to a successful spacer acquisition [16]. Using this information we can estimate $\mu_r$.

In any given infected cell, assuming viral genome abundance increases exponentially with time (linear viral production makes little difference, see section below), virions are produced at a rate $\gamma \log \beta$. We want the total "viral-replicon-hours" per infection during which a cell can obtain a spacer:

$$\eta = \int_0^\tau t e^{\gamma t \log \beta} \, dt = \frac{\beta^{\gamma\tau}(\gamma\tau \log \beta - 1) + 1}{\gamma^2 \log^2 \beta} \tag{13}$$

where $0 < \tau \leq 0.75$ hours is the time-window during infection where spacer acquisition can occur. We can then convert between a per-infection and per-replicon-hour growth rate:

$$\mu_r = \frac{10^{-7}}{\eta}. \tag{14}$$

In the system we are considering, spacer acquisition occurs early on in infection [16]. Thus, in this model system, for $\tau = 10\,\text{min}$, we can estimate that $\mu_r$ is approximately $4 \times 10^{-6}$. If the window in which infection occurs is shorter, then our estimate of $\mu_r$ will be higher (e.g., for $\tau = 5\,\text{min}$ we estimate that $\mu_r \approx 2 \times 10^{-5}$) and if it is longer than our estimate of $\mu_r$ will be lower (e.g., for $\tau = 20\,\text{min}$ we estimate that $\mu_r \approx 4 \times 10^{-7}$).

Obviously these values will vary across systems, but the ranges used above are typical of lytic phage in culture (in organisms common in oligotrophic environments, e.g., the open ocean, these parameters can be quite different [24]).

#### Linear Viral Production

Equation 11 above can be modified to reflect linear production of new viruses in the cell:

$$\dot{I}_v = \overbrace{\delta U V}^{\text{Infection}} + \overbrace{\gamma \beta I}^{\text{Reproduction}} - \overbrace{\left(\frac{\gamma\beta}{\xi}\right) I}^{\text{Immunization Window}} - \overbrace{w I_v}^{\text{Washout}} - \overbrace{\mu_r \left(\frac{I_v^2}{I}\right)}^{\text{Immunization}}. \tag{15}$$

In practice, this alteration does not affect our results - as at equilibrium $I_v = 0$ and we start off most of our simulations with a small population of already-immunized host, making the rate of immunization largely irrelevant (since population growth of the immune host occurs more quickly than new immunization of infected host).

Nevertheless, linear production of viruses will affect our estimates of $\mu$, so that if

$$\eta = \int_0^\tau t\gamma\beta \, dt = \frac{\gamma\beta\tau^2}{2} \tag{16}$$

is the total "viral-replicon-hours" per infection during which a cell can obtain a spacer, where $0 < \tau \leq 0.75$ hours is the time-window during infection where spacer acquisition can occur, we can convert between a per-infection and per-replicon-hour growth rate:

$$\mu_r = \frac{10^{-7}}{\eta}. \tag{17}$$

In the system we are considering, spacer acquisition occurs early on in infection [16]. Thus, in this model system, for $\tau = 10\,\mathrm{min}$, we can estimate that $\mu_r$ is approximately $6.8 \times 10^{-8}$. If the window in which infection occurs is shorter, then our estimate of $\mu_r$ will be higher (e.g., for $\tau = 5\,\mathrm{min}$ we estimate that $\mu_r \approx 2.7 \times 10^{-7}$) and if it is longer than our estimate of $\mu$ will be lower (e.g., for $\tau = 20\,\mathrm{min}$ we estimate that $\mu_r \approx 1.7 \times 10^{-8}$). Again, this has no effect on our overall conclusions, except to further emphasize that the effects of autoimmunity are likely to be negligible (since $\mu_r$ is estimated to be lower in this case).

### SText 3 Spacer Acquisition from Defective Phage

Hynes et al. [7] found evidence that in the type II-A CRISPR-Cas systems of *S. thermophilus* the primary substrate for spacer acquisition are defective phage that adsorb to the cell but are incapable of bringing their replication cycles to completion. They suggest that in their system approximately 10% of phage are naturally defective and that the majority of spacers are acquired from these phage. This work suggests an alternate paradigm for spacer acquisition to those presented in S1 Text and in the main text. We modified our model to consider the scenario where spacers are acquired from defective phages ($V_D$) only:

$$\begin{aligned}
\overbrace{\dot{R}}^{\text{Resources}} &= \overbrace{w(r_0 - R)}^{\text{Flow}} - \overbrace{\frac{evR}{z+R}(S+C)}^{\text{Resource Uptake by Cells}} \\
\overbrace{\dot{S}}^{\text{Susceptible}} &= \left(\overbrace{\frac{vR}{z+R}}^{\text{Growth}} - \overbrace{\delta V}^{\text{Infection}} - \overbrace{\psi\delta V_D}^{\text{Immunization}} - \overbrace{w}^{\text{Flow}}\right) S \\
\overbrace{\dot{C}}^{\text{Immune}} &= \left(\overbrace{\frac{vR}{z+R}}^{\text{Growth}} - \overbrace{w}^{\text{Flow}}\right) C + \overbrace{\psi\delta V_D S}^{\text{Immunization}} \\
\overbrace{\dot{I}}^{\text{Infected}} &= \overbrace{\delta S V}^{\text{Infection}} - \overbrace{\gamma I}^{\text{Lysis}} - \overbrace{wI}^{\text{Flow}} \\
\overbrace{\dot{V}}^{\text{Viruses}} &= \overbrace{w(v_0(1-\varepsilon) - V)}^{\text{Flow}} + \overbrace{\beta(1-\varepsilon)\gamma I}^{\text{Lysis}} - \overbrace{\delta(S+C)V}^{\text{Adsorption}} \\
\overbrace{\dot{V_D}}^{\text{Defective Viruses}} &= \overbrace{w(v_0\varepsilon - V)}^{\text{Flow}} + \overbrace{\beta\varepsilon\gamma I}^{\text{Lysis}} - \overbrace{\delta(S+C)V}^{\text{Adsorption}}
\end{aligned} \tag{18}$$

where $\varepsilon$ is the percent of each phage burst that is defective and $\psi = \frac{\mu}{\varepsilon}$ is the per-infection rate of spacer acquisition from defective phage.

For the linear stability analysis in SText 6, we note that this alternative model gives nearly identical results to the models presented in the main text (Fig 2) as well as S1 Text. Briefly, we consider equilibria of the model where $v_0 > 0$ and $\tilde{C} + \tilde{M} > 0$, such that at equilibrium we always have that $\tilde{S} = 0$ (see SText 6). At such an equilibrium, spacer-acquisition dynamics are no longer relevant to the system, so that the equilibria are nearly identical (substituting in $v_0(1-\varepsilon)$ and $\beta(1-\varepsilon)$ for $v_0$ and $\beta$ respectively).

### SText 4 Minimal Lag Model

To model competition between laggy immunity and a surface mutant strategy during an outbreak we can construct a simple six-parameter model by neglecting the immunization and mutation processes, ignoring viral latency and washout, modeling resources implicitly:

$$\begin{aligned}
\overbrace{\dot{S}}^{\text{Susceptible}} &= r\left(1-\tfrac{S+C+M}{K}\right)S - \delta VS \\
\overbrace{\dot{C}}^{\text{Immune}} &= r\left(1-\tfrac{S+C+M}{K}\right)C - \delta VC + \phi L \\
\overbrace{\dot{M}}^{\text{SM}} &= (1-\kappa)r\left(1-\tfrac{S+C+M}{K}\right)M \\
\overbrace{\dot{L}}^{\text{Lagged}} &= \delta VC - \phi L \\
\overbrace{\dot{V}}^{\text{Viruses}} &= \delta\beta SV - \delta(S+C)V
\end{aligned} \tag{19}$$

which can be reduced even further by re-scaling time and population size to give a four parameter model ($K$ and $r$ can be omitted, retained now for ease of interpretation). Qualitatively this model gives similar results to our previous analysis, where immune cells face a strong lag-induced cost during an outbreak (S5 Fig).

### SText 5 Functional Form of CRISPR Downregulation

The results described in the main text for transcriptional upregulation of the CRISPR locus allow cells in an upregulated state ($C_F$) to return to baseline expression levels ($C$) at a constant rate ($\zeta$). Here we consider two alternative scenarios where (1) cells return to baseline at a non-constant rate controlled by the rate of viral encounter, and (2) cells do not return to the baseline state (transcriptional upregulation is permanent, or at least persists for a very long time relative to the dynamics of the system).

For the first scenario we incorporate a non-linear term into the equations for $C_F$:

$$\dot{C}_F = \left(\overbrace{\frac{vR}{z+R}}^{\text{Growth}} - \overbrace{w}^{\text{Flow}}\right) C_F + \overbrace{\mu\delta VS}^{\text{Immunization}} + \overbrace{\phi L}^{\text{Clearance}} \overbrace{- \zeta_1 C_F\left(\frac{C_F}{C_F+\zeta_2 V}\right)}^{\text{Downregulation}} \tag{20}$$

and $C$:

$$\dot{C} = \left( \overbrace{\frac{vR}{z+R}}^{\text{Growth}} - \overbrace{\delta V}^{\text{Infection}} - \overbrace{w}^{\text{Flow}} \right) C + \overbrace{\zeta_1 C_F \left( \frac{C_F}{C_F + \zeta_2 V} \right)}^{\text{Downregulation}} \tag{21}$$

where $\zeta_1$ is the maximum rate of return and $\zeta_2$ is a half-saturation constant which we let be $\zeta_2 = 1 \frac{\text{cells}}{\text{viruses}}$. For ease of comparison we let $\zeta_1 = \zeta$. The intuition behind this functional form is that the return to a baseline transcriptional state will be prevented if there are still many viruses in the environment that are adsorbing to our transcriptionally upregulated immune cells ($C_F$). The equations above will allow upregulated cells to return to baseline transcription following the linear rate ($\zeta_1$) when viruses are absent from the environment, and will not allow cells to return to baseline transcription (i.e., the rate of return approaches zero) when the multiplicity of infection (MOI) is very high ($V \gg C_F$). This addition is somewhat more realistic than a simple linear rate of downregulation, because it accounts for the fact that re-infection of the upregulated cells will likely prolong their upregulated state.

For the second scenario, we remove the downregulation term from the equations for $C_F$:

$$\dot{C}_F = \left( \overbrace{\frac{vR}{z+R}}^{\text{Growth}} - \overbrace{w}^{\text{Flow}} \right) C_F + \overbrace{\mu \delta V S}^{\text{Immunization}} + \overbrace{\phi L}^{\text{Clearance}} \tag{22}$$

and $C$:

$$\dot{C} = \left( \overbrace{\frac{vR}{z+R}}^{\text{Growth}} - \overbrace{\delta V}^{\text{Infection}} - \overbrace{w}^{\text{Flow}} \right) C. \tag{23}$$

In fact, when $C(0) = 0$ cells/mL, we can also remove the equation for $C$, in effect simplifying this system to the case where CRISPR immunity has no lag. This version of the model assumes that once upregulated transcriptionally, the CRISPR system never returns to its baseline transcriptional state, or returns to that state so slowly as to be irrelevant to model dynamics.

### SText 6 Equilibria of Lag Model and Linear Stability Analysis

Consider our lag system (main text, equations 1-4) neglecting the population of susceptibles (which will always be zero at equilibrium if $v_0 > 0$) with a laggy CRISPR-Cas host ($C$) being competed against an SM strain ($M$):

$$\begin{aligned}
\dot{R} &= w(r_0 - R) - \frac{evr}{z+R}(M + C) \\
\dot{V} &= w(v_0 - V) - \delta V C \\
\dot{C} &= \left( \frac{vR}{z+R} - w - \delta V \right) C + \phi L
\end{aligned}$$

$$\begin{aligned}
\dot{L} &= \delta VC - \phi L - wL \\
\dot{M} &= \left(\frac{(1-k)vR}{z+R} - w\right)M.
\end{aligned}$$

We found the equilibria for this model when both host strains are present (tildes indicate equilibrium value):

$$\begin{aligned}
\tilde{R} &= \frac{zw}{v - vk - w} \\
\tilde{V} &= \left(\frac{\phi + w}{\delta w}\right)\left(\frac{v\tilde{R}}{z+\tilde{R}} - w\right) \\
\tilde{C} &= \frac{w(v_0 - \tilde{V})}{\delta \tilde{V}} \\
\tilde{L} &= \frac{\delta \tilde{V}\tilde{C}}{\phi + w} \\
\tilde{M} &= \frac{w(r_0 - \tilde{R})(z + \tilde{R})}{ev\tilde{R}} - \tilde{C}
\end{aligned}$$

when only the CRISPR-encoding strain is present:

$$\begin{aligned}
\tilde{V} &= \frac{wv_0}{w + \delta\tilde{C}} \\
\tilde{C} &= \frac{w(r_0 - \tilde{R})(z + \tilde{R})}{ev\tilde{R}} \\
\tilde{L} &= \frac{\delta \tilde{V}\tilde{C}}{\phi + w} \\
\tilde{M} &= 0
\end{aligned}$$

and solving for $\tilde{R}$ (using Mathematica 12 [8]):

$$\frac{wv_0 ev\tilde{R}}{wev\tilde{R} + \delta w(r_0 - \tilde{R})(z + \tilde{R})} = \left(\frac{\phi + w}{\delta w}\right)\left(\frac{v\tilde{R}}{z+\tilde{R}} - w\right)$$

when only the SM strain is present:

$$\begin{aligned}
\tilde{R} &= \frac{zw}{v - vk - w} \\
\tilde{V} &= v_0 \\
\tilde{C} &= 0 \\
\tilde{L} &= 0 \\
\tilde{M} &= \frac{w(r_0 - \tilde{R})(z + \tilde{R})}{ev\tilde{R}}
\end{aligned}$$

and in the absence of host:

$$\begin{aligned}
\tilde{R} &= r_0 \\
\tilde{V} &= v_0 \\
\tilde{C} &= 0 \\
\tilde{L} &= 0 \\
\tilde{M} &= 0.
\end{aligned}$$

In order to assess the stability of our equilibria we linearized our system around each equilibrium point. We found the Jacobian:

$$J = \begin{bmatrix}
-w - \frac{evz(M+C)}{(z+R)^2} & 0 & \frac{-evR}{z+R} & 0 & \frac{-evR}{z+R} \\
0 & -(w+\delta C) & -\delta V & 0 & 0 \\
\left(\frac{vz}{(z+R)^2}\right) C & -\delta C & \left(\frac{vR}{z+R}\right) - w - \delta V & \phi & 0 \\
0 & \delta C & \delta V & -(\phi + w) & 0 \\
\left(\frac{(1-\kappa)vz}{(z+R)^2}\right) M & 0 & 0 & 0 & \frac{(1-\kappa)vR}{z+R} - w
\end{bmatrix}$$

and calculated the eigenvalues of this matrix for each equilibrium point. When all eigenvalues have negative real parts, the equilibrium is stable. If at least one eigenvalue has a real part greater than zero, the equilibrium is unstable (for eigenvalues with real parts equal to zero no determination can immediately be made - though this situation did not come up in our analyses. For further information find standard procedures for linear stability analysis in any text on ordinary differential equations).

### SText 7 Implicit Resource Dynamics

Consider a modified version of our lag system (main text equations 1-4) with implicit (logistic growth) rather than explicit resource dynamics:

$$\begin{aligned}
\overbrace{\dot{S}}^{\text{Susceptible}} &= r(1 - \tfrac{S+C+M}{K})S - \delta VS - wS \\
\overbrace{\dot{C}}^{\text{Immune}} &= r(1 - \tfrac{S+C+M}{K})C + \mu\delta VS - \delta VC + \phi L - wC \\
\overbrace{\dot{I}}^{\text{Infected}} &= (1-\mu)\delta VS - \gamma I - wI \\
\overbrace{\dot{M}}^{\text{SM}} &= (1-\kappa) r(1 - \tfrac{S+C+M}{K})M - wM \\
\overbrace{\dot{L}}^{\text{Lagged}} &= \delta VC - \phi L - wL \\
\overbrace{\dot{V}}^{\text{Viruses}} &= \delta\beta SV - \delta(S+C)V + w(v_0 - V)
\end{aligned} \tag{24}$$

Neglecting the population of susceptibles (which will always be zero at equilibrium if $v_0 > 0$) with a laggy CRISPR-Cas host ($C$) being competed against an SM strain ($M$) we found the equilibria when only CRISPR host are present:

$$\begin{aligned}\tilde{V} &= \frac{wv_0}{\delta\tilde{C} + w}\\ \tilde{L} &= \frac{\delta\tilde{V}\tilde{C}}{\phi + w}\\ \tilde{M} &= 0\end{aligned}$$

and solving for $\tilde{C}$ (using Mathematica 12 [8]):

$$\left(\frac{\delta wv_0}{\delta\tilde{C} + w}\right)\left(\frac{1}{\phi + w}\right) = \left(\frac{1}{w}\right)\left(r\left(1 - \frac{\tilde{C}}{K}\right) - w\right)$$

when only SM host are present:

$$\begin{aligned}\tilde{V} &= v_0\\ \tilde{C} &= 0\\ \tilde{L} &= 0\\ \tilde{M} &= K\left(1 - \frac{w}{(1-\kappa)r}\right)\end{aligned}$$

and when no host are present:

$$\begin{aligned}\tilde{V} &= v_0\\ \tilde{C} &= 0\\ \tilde{L} &= 0\\ \tilde{M} &= 0.\end{aligned}$$

In order to assess the stability of our equilibria we linearized our system around each equilibrium point. We found the Jacobian:

$$J = \begin{bmatrix} r\left(1 - \frac{2C+M}{K}\right) - \delta V - w & \frac{-rC}{K} & \phi & 1 - \delta C\\ \frac{-r(1-\kappa)M}{K} & (1-\kappa)r\left(1 - \frac{C+2M}{K}\right) - w & 0 & 0\\ \delta V & 0 & -(\phi + w) & \delta C\\ -\delta V & 0 & 0 & -\delta C - w \end{bmatrix}$$

and calculated the eigenvalues of this matrix for each equilibrium point. When all eigenvalues have negative real parts, the equilibrium is stable. If at least one eigenvalue has a real part greater than zero, the equilibrium is unstable (for eigenvalues with real parts equal to zero no determination can immediately be made - though this situation did not come up in our analyses. For further information find standard procedures for linear stability analysis in any text on ordinary differential equations).

Similar to our lag model in the main text (see Figure 2), this model with implicit resource dynamics showed a region of bistability, where both CRISPR-only and SM-only states were stable, and which state the system ultimately would fall into depends on the initial conditions (S8 Figure). We hypothesized that this bistability was a byproduct of the degradative nature of CRISPR immunity. That is, the CRISPR immune strain is able to rapidly clear viruses from the environment (since it is a DNA-degrading intracellular immune system), which can in turn facilitate it's establishment in these cases by removing the source of lag (*i.e.*, viruses). To demonstrate this point, we consider a modified scenario where CRISPR does not clear viruses from the environment:

$$\begin{aligned}
\overbrace{\dot{S}}^{\text{Susceptible}} &= r(1-\tfrac{S+C+M}{K})S - \delta VS - wS \\
\overbrace{\dot{C}}^{\text{Immune}} &= r(1-\tfrac{S+C+M}{K})C + \mu\delta VS - \delta VC + \phi L - wC \\
\overbrace{\dot{I}}^{\text{Infected}} &= (1-\mu)\delta VS - \gamma I - wI \\
\overbrace{\dot{M}}^{\text{SM}} &= (1-\kappa)r(1-\tfrac{S+C+M}{K})M - wM \\
\overbrace{\dot{L}}^{\text{Lagged}} &= \delta VC - \phi L - wL \\
\overbrace{\dot{V}}^{\text{Viruses}} &= \delta\beta SV - \delta SV + w(v_0 - V)
\end{aligned} \tag{25}$$

This scenario is not biologically realistic, but it is conceptually informative. In this case the no-host and SM-only equilibria are identical, but the CRISPR only equilibrium becomes:

$$\begin{aligned}
\tilde{V} &= v_0 \\
\tilde{C} &= K\left(1-\left(\frac{w}{r}\right)\left(\frac{\delta V_0}{\phi+w}+1\right)\right) \\
\tilde{L} &= \frac{\delta\tilde{V}\tilde{C}}{\phi+w} \\
\tilde{M} &= 0.
\end{aligned}$$

Finding the Jacobian of this system, and performing linear stability analysis as above, we find that when CRISPR does not clear viruses from the environment the region of bistability disappears (S9 Figure).

[8] Wolfram Research, Inc. Mathematica, Version 12.1. Champaign, IL, 2020.

[23] Jake L Weissman, Arlin Stoltzfus, Edze R Westra, and Philip LF Johnson. Avoidance of self during CRISPR immunization. *Trends in Microbiology*, 2020.

[24] Joshua S. Weitz. *Quantitative viral ecology: dynamics of viruses and their microbial hosts*. Princeton University Press, January 2016.

[25] Joshua S Weitz, Charles A Stock, Steven W Wilhelm, Lydia Bourouiba, Maureen L Coleman, Alison Buchan, Michael J Follows, Jed A Fuhrman, Luis F Jover, Jay T Lennon, et al. A multitrophic model to quantify the effects of marine viruses on microbial food webs and ecosystem processes. *The ISME journal*, 9(6):1352–1364, 2015.

[26] Edze R. Westra, Stineke van Houte, Sam Oyesiku-Blakemore, Ben Makin, Jenny M. Broniewski, Alex Best, Joseph Bondy-Denomy, Alan Davidson, Mike Boots, and Angus Buckling. Parasite exposure drives selective evolution of constitutive versus inducible defense. *Curr Biol*, 25(8):1043–1049, April 2015.

[27] X Sunney Xie, Paul J Choi, Gene-Wei Li, Nam Ki Lee, and Giuseppe Lia. Single-molecule approach to molecular biology in living bacterial cells. *Annu. Rev. Biophys.*, 37:417–444, 2008.

| Parameter | Definition | Value |
|---|---|---|
| $\alpha$ | Max. Rate of Genome Replication | $\sim 0.4$ genomes/hr (theoretical max for a genome size 5-6mb) |
| $\mu_r$ | Per-Replicon Spacer Acquisition Rate | $6.8 \times 10^{-8}$ per replicon per hour section §SText 2 |
| $\psi$ | Probability of Spacer Acquisition from a Defective Phage | $\frac{\mu}{\varepsilon}$ |
| $\varepsilon$ | Proportion of Phage that are Defective | 0.01[7] |
| $\xi$ | Immunization Window | 10 minutes |

Table S1: Parameter definitions for supplemental models

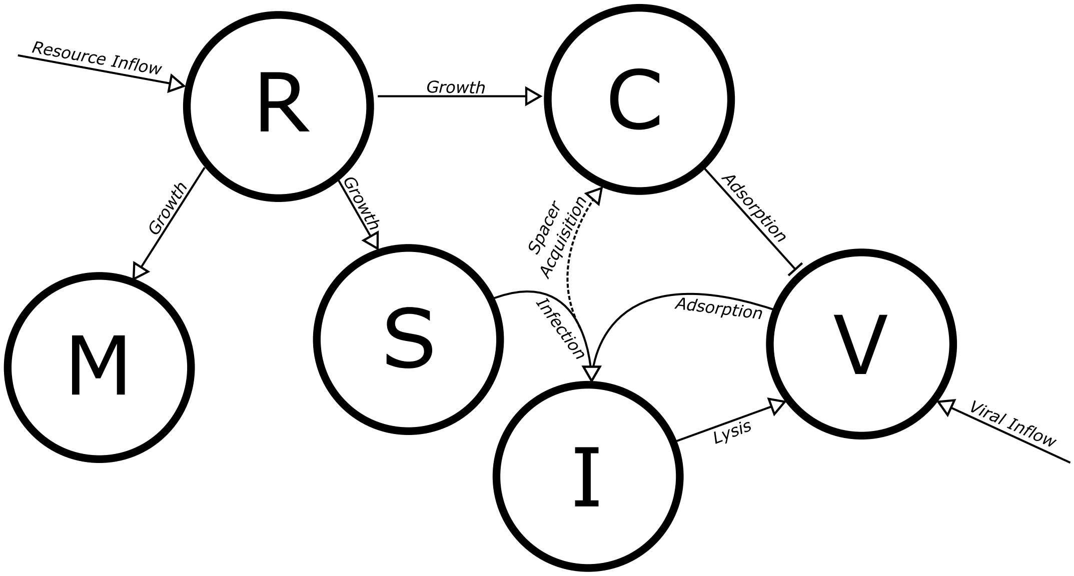


Figure S1: Diagram of base model without lag.

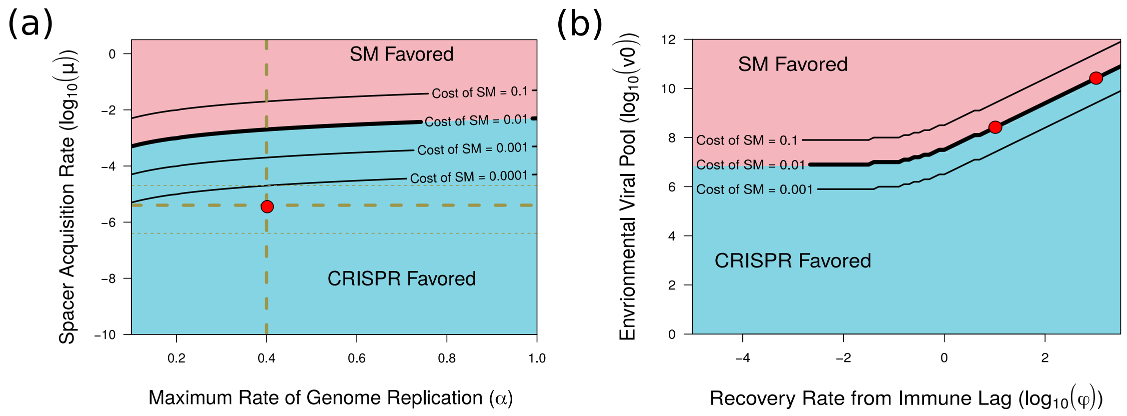


Figure S2: Immune lag, but not autoimmunity, can lead to invasion of an SM strategy into a resident CRISPR-encoding population (using replicon model in section §SText 2). Colors and thick black line denote regions of parameter space where invasion can (pink) or cannot (blue) occur when SM is associated with a 1% growth cost. Thin black lines show how this boundary changes with different growth costs. Dashed lines and red circles in (a) indicate best estimates for parameter values ($\mu_r = 4 \times 10^{-6}$ and $\alpha = 0.4$; thin lines at $\mu_r = 2 \times 10^{-5}$ and $\mu_r = 4 \times 10^{-7}$). Red circles in (b) denote approximate phage concentrations and inferred $\phi$ where the cost of CRISPR-Cas increases sharply in competition experiments ($\phi = 10$, $\phi = 10^3$). (a) Shows the invasion condition from equation S8. (b) Shows the numerical result from our replicon model with a small initial SM population ($M(0) = 100$), large initial CRISPR-encoding population ($C(0) = 10^8$), and no susceptibles (solved at $t = 10^7$ hours to approximate equilibrium).

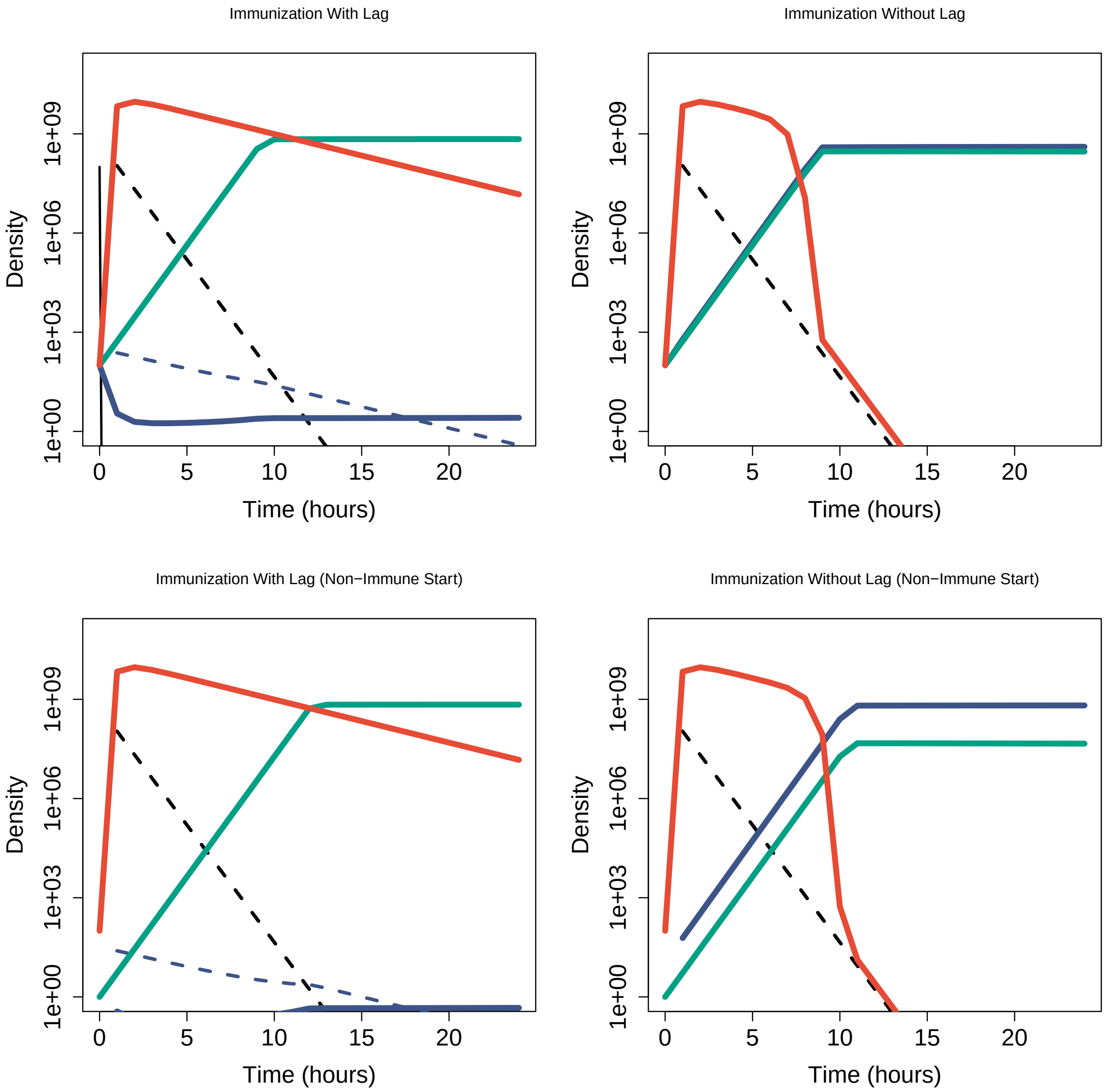


Figure S3: Immune lag is costly during an outbreak of novel phage (alternative model of spacer acquisition from defective phage). Top panels depict system initialized with a dense population of susceptible host ($S = 10^8$) and small populations of SM and CRISPR-immune host and virus ($C = M = 100$). Bottom panels depict system without substantial immune/resistant host at $t = 0$ ($C = 0$, $M = 1$).

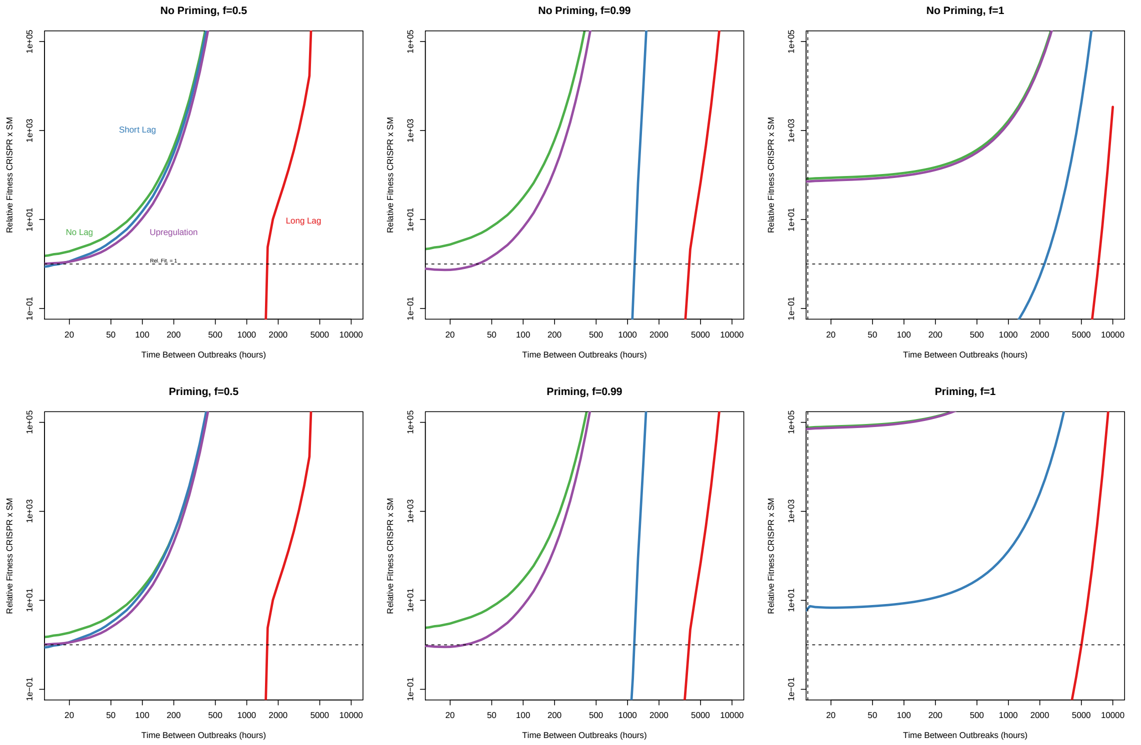


Figure S4: Priming can help CRISPR-defended host when outbreaks of novel virus affect the entire host population, but is only marginally helpful if even a small fraction of host remain unaffected by the outbreak. We let primed adaptation occur at 1000 times the rate of naive adaptation, assuming that the novel virus at each outbreak was not targeted by the host CRISPR-array at the outset but that a partial match would lead to rapid primed adaptation. In practice, this simply means multiplying $\mu$ by a factor of 1000 in our simulations. When $f = 1$, we let $M = 1$ at the start of each outbreak, under the assumption that a rare preexisting SM was already present in the population, in order to avoid modeling mutation dynamics (given large host population sizes at equilibrium). Otherwise results are the same as in Fig 4d. Since host will not reach maximum density until $\sim 12$ hours after an outbreak, we do not consider outbreaks more frequent than this interval.

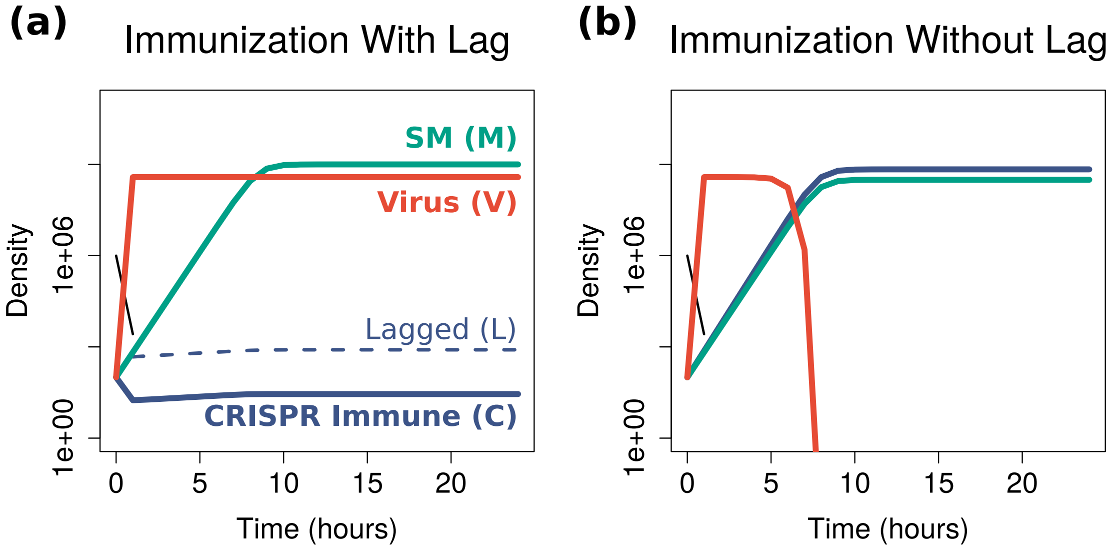


Figure S5: Immune lag is costly during an outbreak of novel phage (minimal model of immune lag; $r = 2$, $K = 10^9$, $\kappa = 0.01$, $\delta = 10^{-7}$, $\beta = 80$, $\phi = 10$). System initialized with a dense population of susceptible host ($10^8$) and small populations of SM and CRISPR-immune host and virus (100).

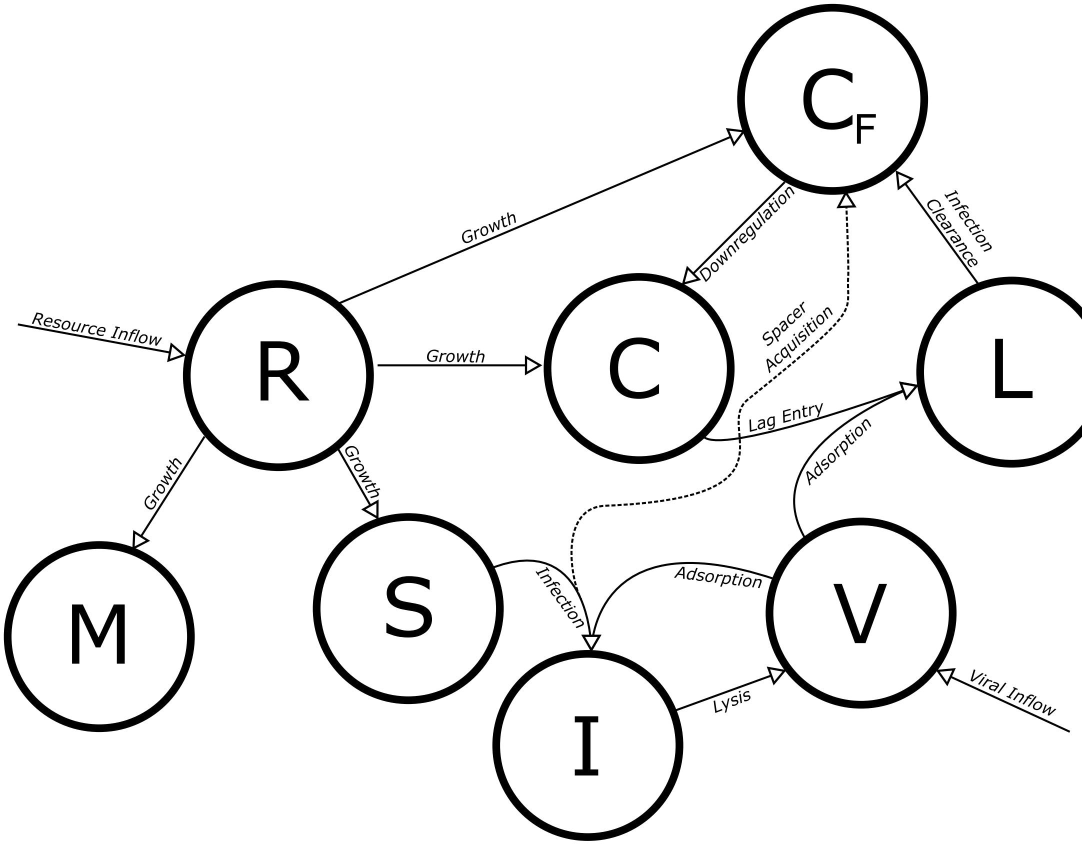


Figure S6: Diagram of lag model with transcriptional upregulation of the *cas* locus.

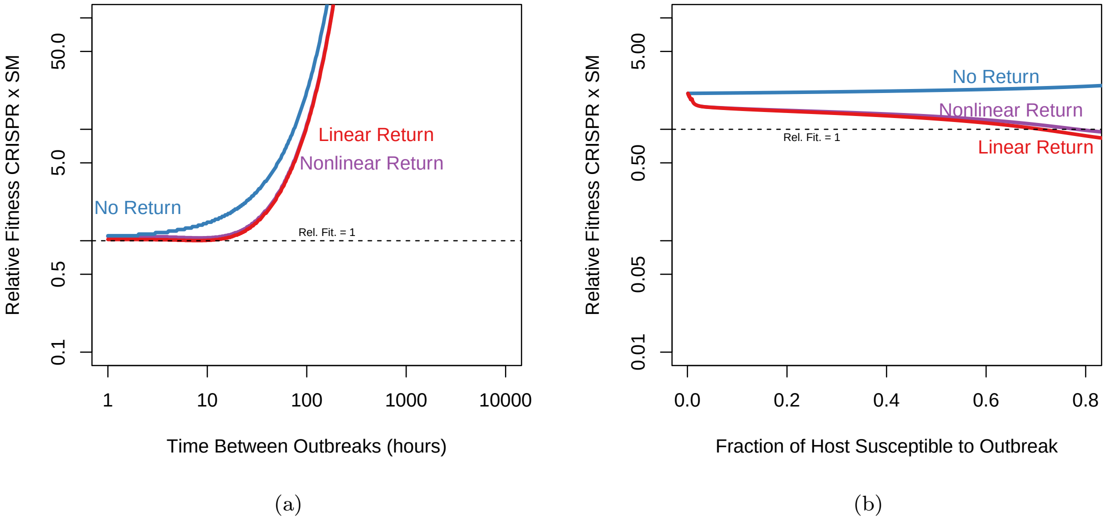


Figure S7: Different functional forms for the return of transcriptionally upregulated cells to baseline transcription do not change overall model results. In all cases, upregulation of the CRISPR array can mitigate the effects of lag (compare with Fig 4). See S5 Text for a description of the different functional forms for the rate of return to baseline visualized here.

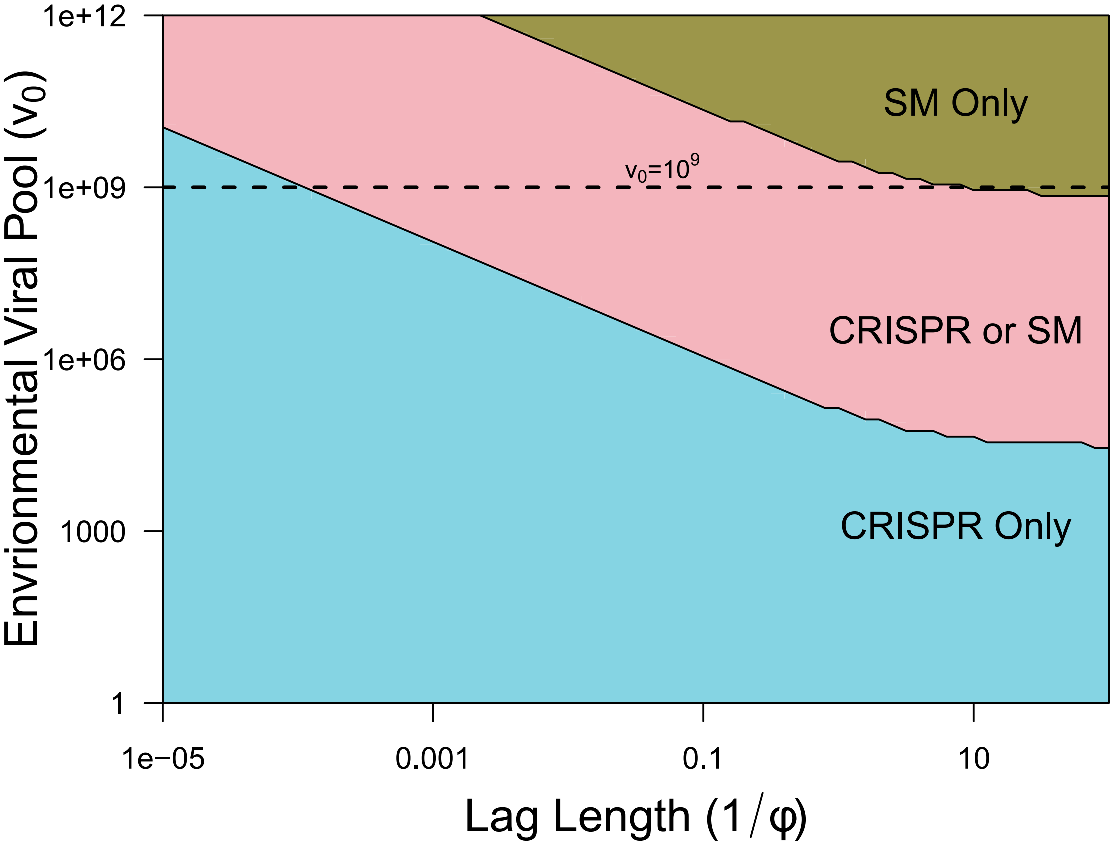


Figure S8: Stable equilibria of lag model with implicit resource dynamics.

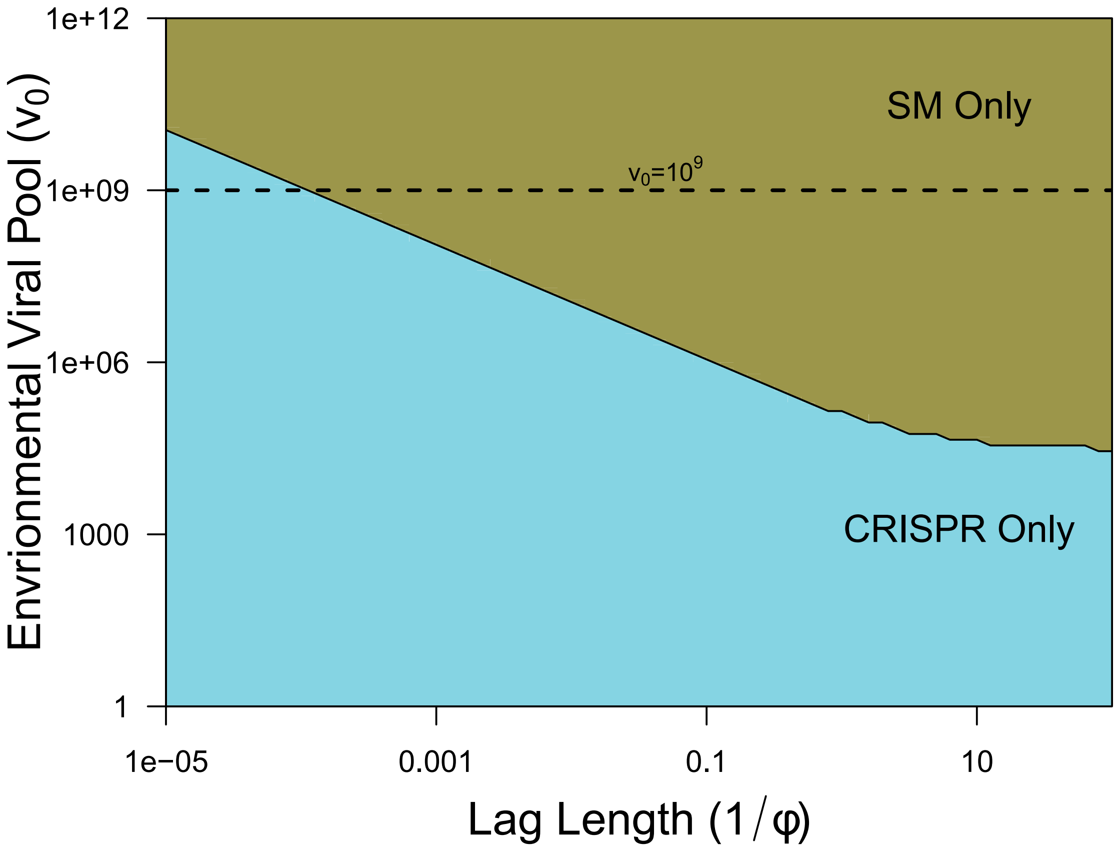


Figure S9: Stable equilibria of lag model with implicit resource dynamics when CRISPR is unable to clear viruses from the environment.

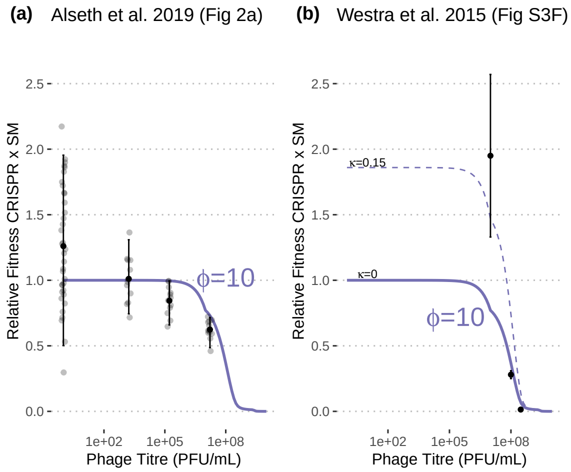


Figure S10: A strongly inducible cost of CRISPR-Cas immunity at high viral titre is consistent with "laggy" CRISPR-Cas immunity. Results of similar previous experiments with the same strains (b-c; [2, 26]) are similar to our own, though they suggest a much longer lag period (fit for model with $\phi = 10$ shown) and disagree on the baseline relative fitness of the CRISPR-immune verus SM strains in the absence of virus ((b) implies a costly mutation in the SM strain).

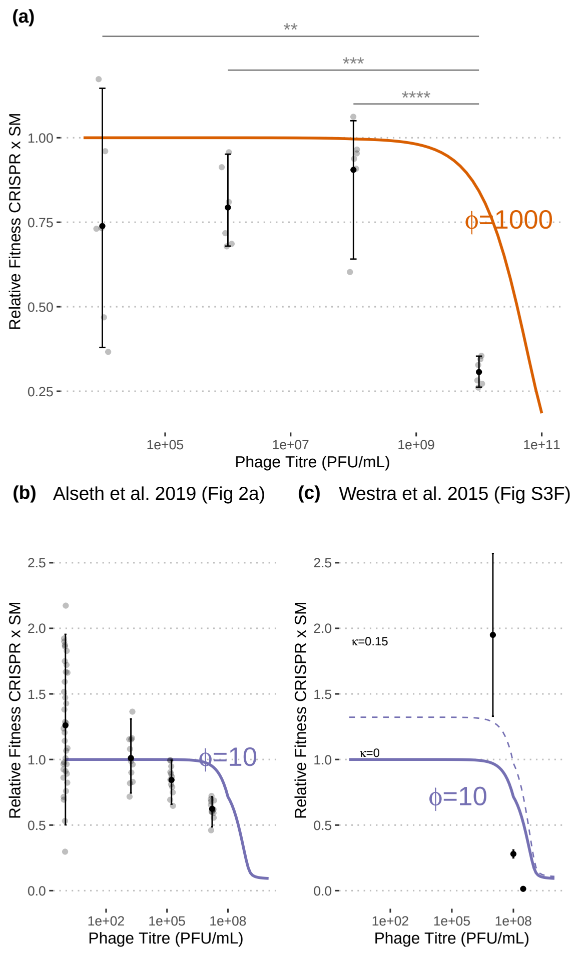


Figure S11: Experimental data as compared to simulations starting with a 10-fold larger initial concentration of susceptible cells ($10^8$CFU/mL).

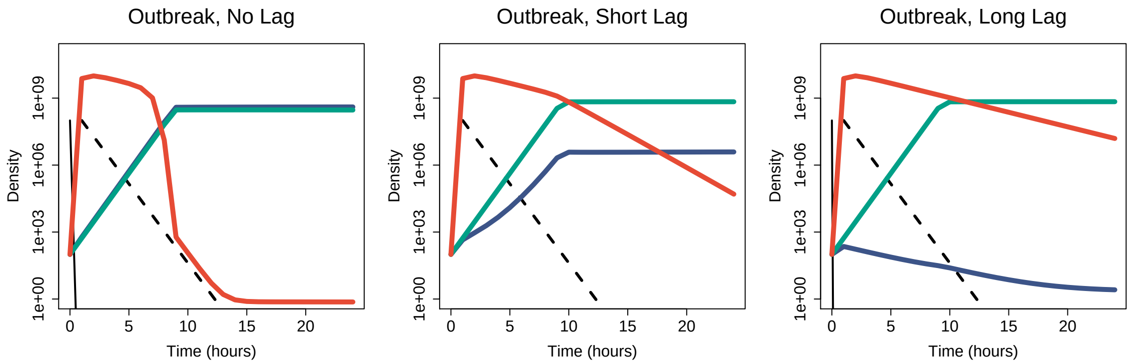


Figure S12: Immune lag prevents the expansion of a CRISPR strategy during an outbreak of novel phage. Same model solutions as in Figure 2 panels a-c, but with populations of lagged and un-lagged CRISPR strains shown combined (purple line).

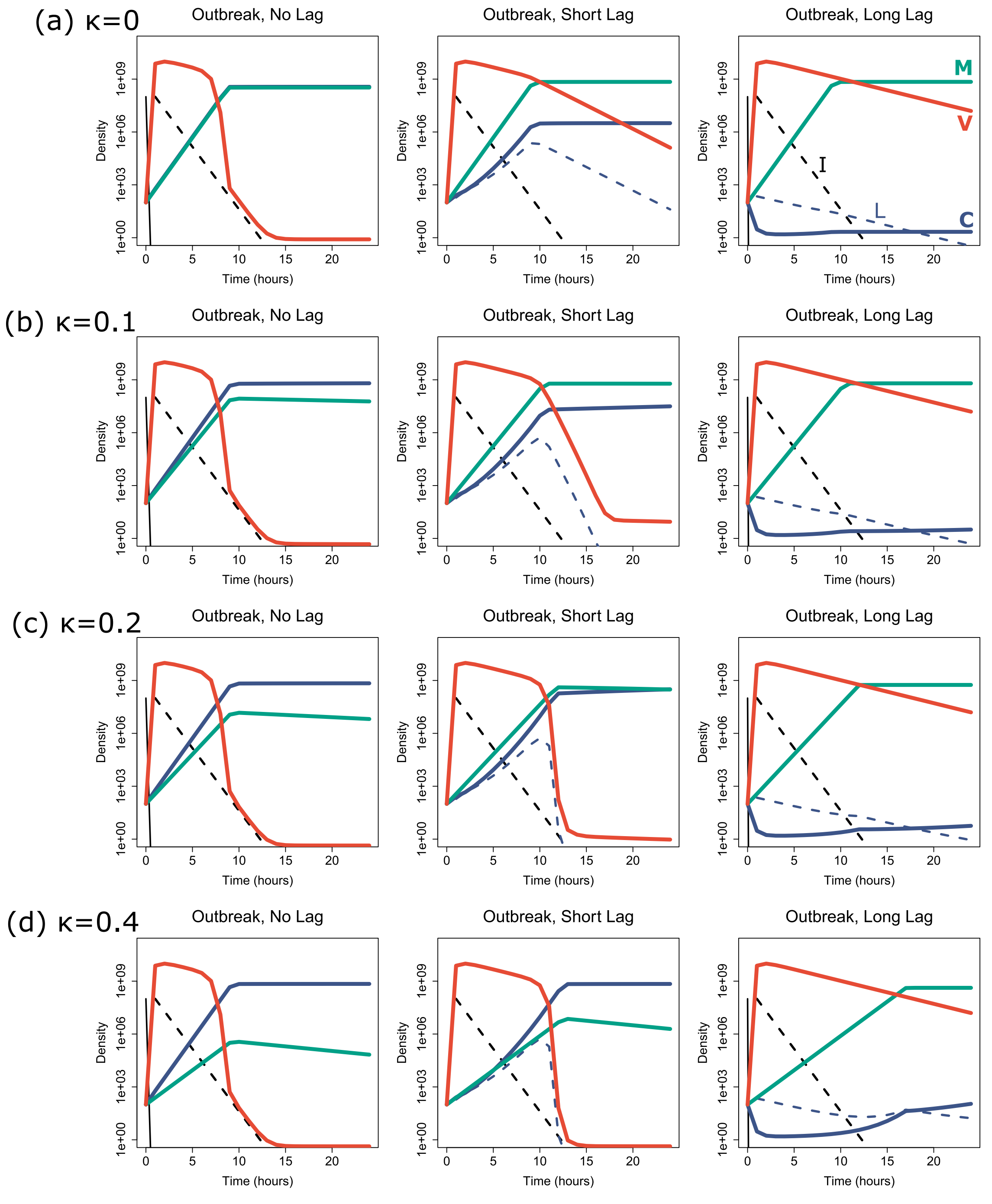


Figure S13: Immune lag prevents the expansion of a CRISPR strategy during an outbreak of novel phage even with a very costly SM strain. With a short lag ($\phi = 10^3$), even with a 20% growth cost the SM strain still outcompetes the CRISPR strain during the early outbreak, and if the lag is long ($\phi = 10$) then the SM strain can outcompete the CRISPR strain even when it has an extremely high 40% growth cost. System initialized with a dense population of susceptible host ($10^8$) and small populations of SM and CRISPR-immune host and virus (100).

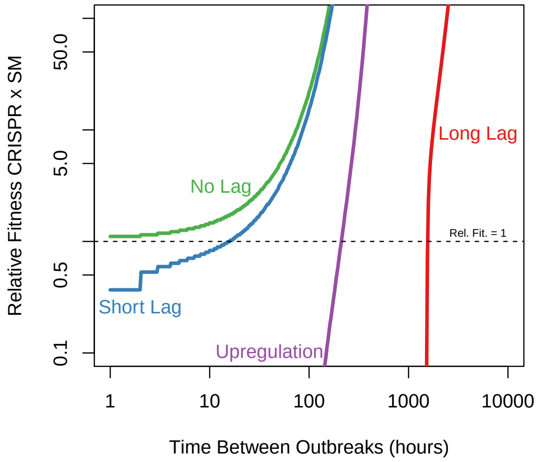


(a)

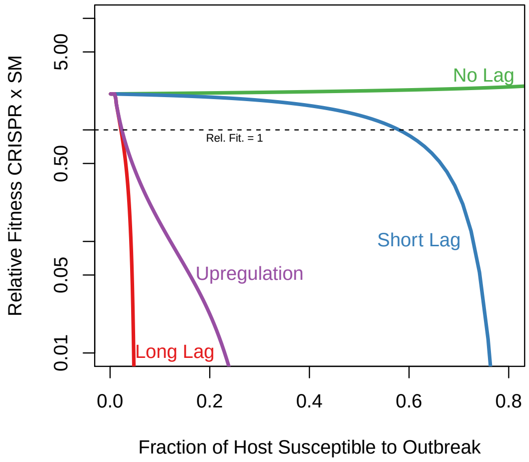


(b)

Figure S14: If the return to normal expression levels from an upregulated-CRISPR state is fast ($\zeta = 10$), then the beneficial effects of upregulation during an outbreak are diminished (though still apparent).
